# Temporal instability weakens entorhinal grid-like coding in older adults

**DOI:** 10.64898/2026.10.15.759328

**Authors:** Vladislava Segen, Gerard Callau Navarro, Philip Bahrd, Tuğçe Belge Bernard, Julia-Isabel Schinke, Thomas Wolbers

**Author notes:** Correspondence and lead contact: Vladislava Segen (*).

## Abstract

Entorhinal grid cell representations are thought to support spatial navigation and episodic memory, functions that decline with age. Their putative fMRI correlate, the hexadirectional signal, varies markedly among older adults, but the source of this variability is unknown. Using virtual navigation and fMRI in 85 older adults, we tested whether signal strength was related to the stability of hexadirectional signals. Hexadirectional orientation stability, the cross-run reproducibility of direction-specific activity patterns, was associated with hexadirectional signal strength in the posterior-medial-entorhinal-cortex (pmEC). In left-pmEC, higher orientation stability also predicted better object-location memory. In contrast, age and local structural measures did not account for individual differences in either measure. Finally, retrosplenial absolute-orientation stability was associated with left-pmEC orientation stability, and a mediation analysis supported an indirect association with hexadirectional signals through pmEC orientation stability. Together, these findings suggest that weaker entorhinal hexadirectional signal in aging may reflect unstable, rather than absent, entorhinal grid-cell-like representations.

## INTRODUCTION

Normal aging is typically associated with declines in episodic memory and spatial navigation, but these changes often vary considerably across individuals: some older adults retain high levels of performance, whereas others show marked impairment^1–4^. Both functions depend in part on medial temporal lobe circuits, including entorhinal grid cells, which provide a periodic, hexagonally organized spatial code that supports self-location during navigation^5^. In humans, grid signal representations can be indexed with fMRI through a characteristic hexadirectional modulation of activity by movement direction^6^. On average, older adults show weaker entorhinal grid-cell-like signals than young adults, yet some express grid-cell-like signals comparable to those of young adults^7^. Thus, grid-like coding is not uniformly lost in later life, but what accounts for its marked individual variability remains unknown.

Animal work suggests that age-related variability in spatial coding need not reflect disappearance of the underlying neural code. Hippocampal place cells remain spatially tuned in aged rodents, but their maps are less stable over repeated experience, show increased remapping, and can become misaligned with external cues^8,9^. This disruption appears to extend to the medial entorhinal cortex: despite preserved grid-cell density, aged animals show reduced grid-cell stability and less reliable remapping in response to environmental change^10^. Thus, a critical unresolved question is not simply whether grid coding persists in human aging, but what determines how reliably it is expressed.

Several forms of representational instability could weaken a macroscopic grid-cell-like signal. Grid orientation may drift over time, preferred hexadirectional orientations may become less coherent across entorhinal voxels, or the multivoxel geometry of direction-related activity may fail to reproduce over time^11,12^. These possibilities are distinct because the fMRI grid-cell-like signal depends not only on the presence of spatially periodic responses, but also on the precision, alignment, and temporal consistency with which activity is organized across the population. Convergent computational evidence indicates that hexadirectional population activity is weakened when directional tuning becomes broader or less precisely aligned with the underlying grid axes^13^. At the fMRI level, reduced coherence of preferred hexadirectional orientations could therefore attenuate the aggregate signal even if local hexadirectional responses remain present.

A second knowledge gap pertains to the source of this functional instability. One possibility is that unreliable orientation and heading information compromises the stability of grid representations. In rodents, disruption of the anterior thalamic head-direction network markedly impairs the spatial periodicity of entorhinal and parasubicular grid cells^14^. In humans, orientation-related signals have been identified in retrosplenial and entorhinal-subicular regions^15–18^, and healthy aging may compromise the fidelity of this system: older adults show greater angular errors and increasing heading uncertainty over delays, effects reproduced by synaptic noise and modest neuronal loss in a ring-attractor model^19^. Less stable orientation information could thus weaken the directional anchoring of entorhinal grid codes over time. Whether instability in orientation-related representations contributes to individual differences in grid-cell-like coding among older adults also remains unknown.

To address these knowledge gaps, we tested whether individual differences in grid-cell-like signal in the pmEC of healthy older adults were explained by functional instability of the underlying representation. We then asked which form of instability was most strongly associated with grid-cell-like signal and whether unstable orientation-related coding in retrosplenial cortex and presubiculum was associated with temporal instability in pmEC. To preview our findings, grid-cell-like signal was most consistently associated with hexadirectional orientation stability, which also predicted object-location memory. In addition, retrosplenial absolute-orientation stability was associated with pmEC orientation stability and showed an indirect association with grid-cell-like signal through this measure. Together, these findings identify temporal instability in grid-relative orientation patterns as a key correlate of individual differences in pmEC hexadirectional signal in older adults.

## RESULTS

### Hexadirectional signal varies across older adults independently of age

To investigate what drives variability in grid-cell-like coding in older age, we analyzed data from 85 cognitively unimpaired older adults who completed a virtual-navigation task during fMRI. Participants passively moved through a square virtual environment while encoding the locations of five target objects, followed by a three-alternative forced-choice object-location test. In total, participants completed two runs, consisting of the same environment each with a new set of objects in new locations (Figure 1A).

**Figure 1.**
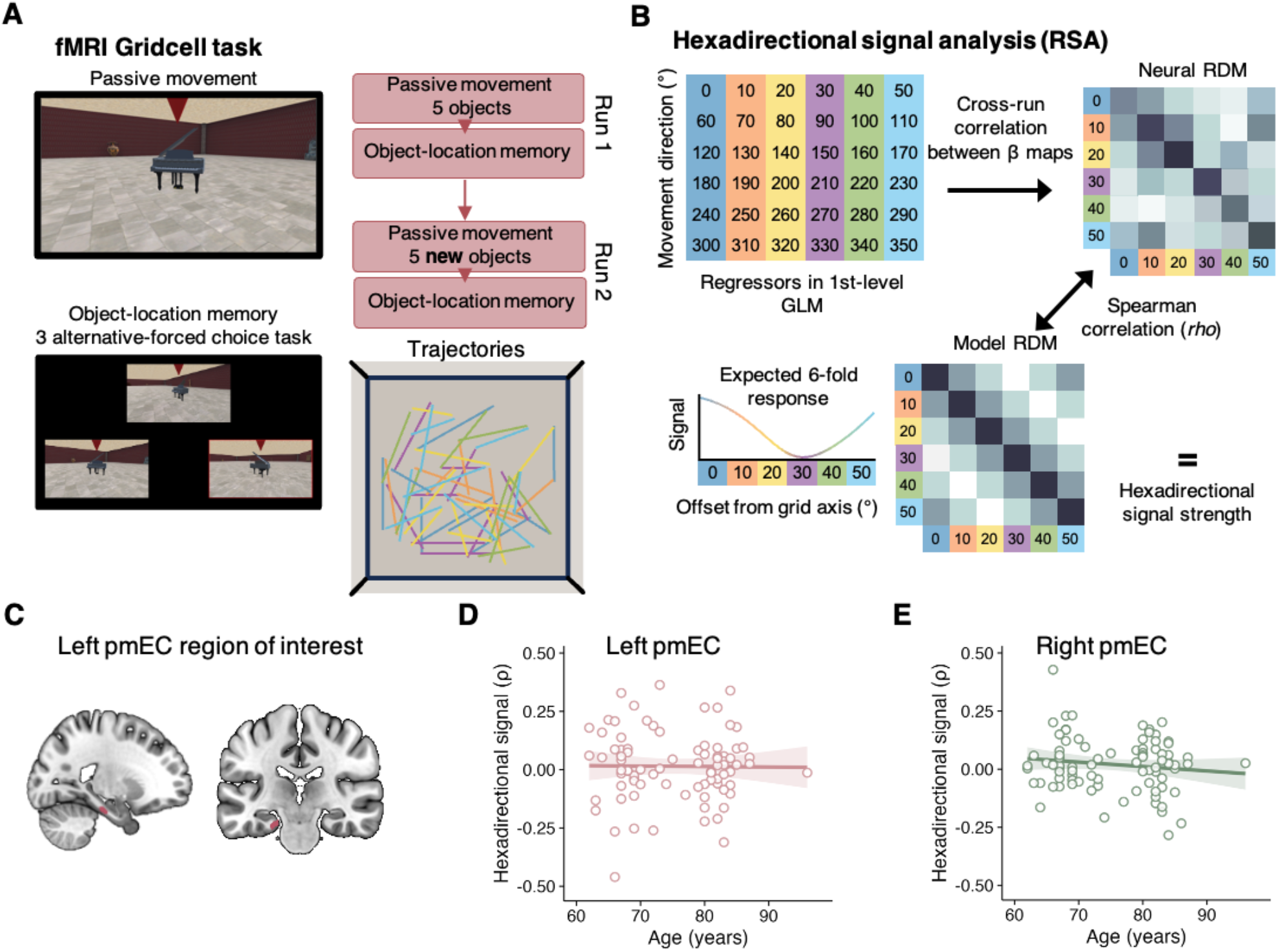
(A) Participants performed two runs of a virtual navigation fMRI task, where they were passively moved through a square environment while encoding five object locations; each run was followed by a three-alternative forced-choice object-location memory test. (B) Hexadirectional signal was estimated with representational similarity analysis (RSA). Direction-specific beta patterns from independent runs were used to form a cross-run neural representational dissimilarity matrix (RDM), which was compared with a hexadiractional model RDM. The Spearman correlation between the two RDMs provided the participant-specific estimate of hexadirectional signal strength. (C) Example of left posterior-medial entorhinal cortex (pmEC) regions of interest used for the RSA. (D,E) Associations between age and hexadirectional signal in left (D) and right (E) pmEC. Each point represents one participant with shaded bands representing 95% confidence intervals.

We quantified hexadirectional signal, a common proxy for population-level grid-cell-like code in humans^6^, using multivariate representational similarity analysis. To index hexadirectional signal in left and right pmEC, we assessed whether cross-run multivoxel activity patterns from six movement-direction bins, defined in 60°-folded space, showed the similarity structure predicted by a hexadirectional model. The Spearman correlation between each participant’s neural and model representational dissimilarity matrices served as the estimate of hexadirectional signal (Figure 1B; see Methods).

Consistent with previous findings^7^, pmEC hexadirectional signal showed marked inter-individual variability (Figure 1D/E). Some showed clear positive hexadirectional modulation, whereas others showed weak or absent hexadirectional signal. This variability was not explained by chronological age in the left, (β = −0.010, 95% CrI [−0.225, 0.204], BF₀₁ = 4.633) or right pmEC, (β = −0.126, 95% CrI [−0.339, 0.085], BF₀₁ = 2.329). Thus, within this older-adult sample, chronological age alone did not account for individual differences in pmEC hexadirectional signal.

### Stable orientation coding supports pmEC hexadirectional signal and object-location accuracy

Rodent work suggests that aging may weaken medial entorhinal coding by reducing the stability of spatial representations rather than eliminating them^10^. To capture complementary aspects of representational reliability in fMRI, we quantified three stability measures separately in each pmEC hemisphere (Figure 2A): **hexadirectional orientation stability**, the reproducibility of matching grid-relative multivoxel patterns across runs; **hexadirectional geometry stability**, the reproducibility of relationships among direction-specific patterns across runs; and **hexadirectional spatial consistency**, the consistency of orientation-specific activity across pmEC voxels. These measures differ from conventional fMRI grid signal stability metrics^7,20^,which assess the alignment or reproducibility of voxelwise preferred six-fold orientations, our measures retain direction-specific responses and weight all grid-relative directions equally. Temporal orientation stability captures the reproducibility of multivoxel patterns across runs, whereas geometry stability captures the reproducibility of their pairwise similarity structure. Spatial consistency indexes response-amplitude dispersion across voxels for each grid-relative direction. These complementary measures follow the representational-similarity framework used to estimate the hexadirectional signal. Importantly, these metrics were weakly correlated and showed minimal collinearity (maximum |**r**| = 0.172; maximum VIF = 1.115), suggesting that they captured distinct properties that could make separable contributions to pmEC hexadirectional signal.

**Figure 2.**
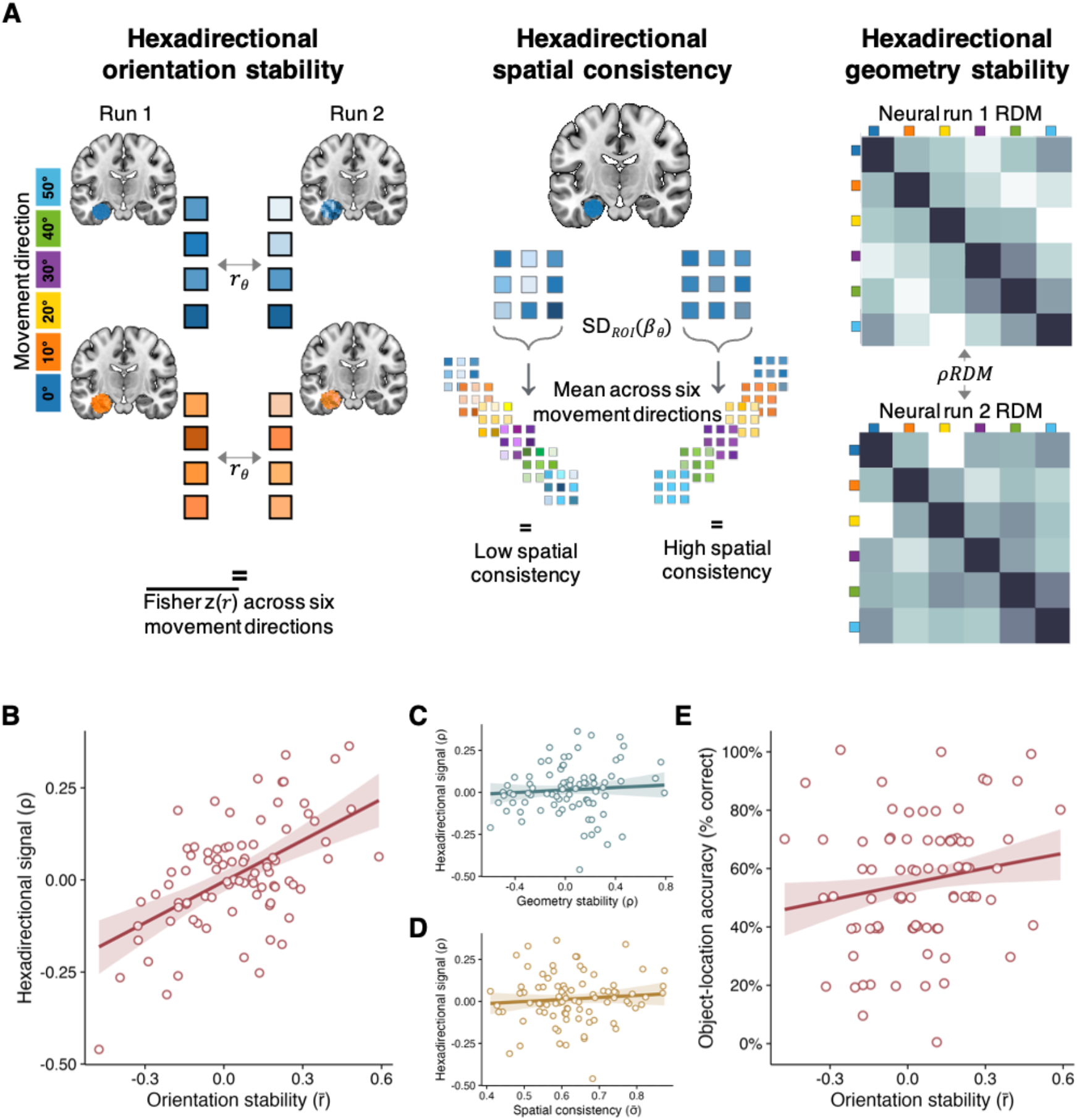
(A) Schematic of the three stability measures: **hexadirectional orientation stability**, the mean Fisher z-transformed correlation between matching grid-relative multivoxel patterns across runs; **hexadirectional spatial consistency**, the across-voxel standard deviation of direction-specific beta estimates, averaged across movement directions and runs; and **temporal hexadirectional geometry stability**, the correlation between the run-specific neural representational dissimilarity matrices (RDMs). (B–D) Joint-model associations of left-pmEC hexadirectional signal with orientation stability (B), geometry stability (C), and spatial consistency (D), adjusting for age and the other stability measures. Fitted lines show posterior predictions with non-plotted predictors held at their sample means. (E) Association between left-pmEC hexadirectional orientation stability and object-location accuracy in an age-adjusted binomial model.

In left-pmEC, in a joint model adjusting for age and the other stability metrics, hexadirectional signal was selectively predicted by hexadirectional orientation stability. Specifically, participants with higher hexadirectional orientation stability showed stronger left-pmEC hexadirectional signal (Figure 2B; β = 0.536, 95% CrI [0.355, 0.718]). By contrast, hexadirectional geometry stability (Figure 2C; β = 0.075, 95% CrI [−0.110, 0.259]) and hexadirectional spatial consistency (Figure2D; β = 0.088, 95% CrI [−0.103, 0.277]) showed no association. Direct posterior comparisons confirmed that the coefficient for hexadirectional orientation stability was larger than that for temporal hexadirectional geometry stability (Δβ = 0.462, 95% CrI [0.186, 0.733]) and than that for hexadirectional spatial consistency (Δβ = 0.448, 95% CrI [0.180, 0.711]).

A parallel analysis in right-pmEC showed a similar association between hexadirectional orientation stability and hexadirectional signal, (β = 0.326, 95% CrI [0.124, 0.529]), while neither hexadirectional geometry stability, (β = 0.179, 95% CrI [−0.016, 0.376]), nor hexadirectional spatial consistency, (β = −0.043, 95% CrI [−0.243, 0.155]), showed a clear association. Direct posterior comparisons indicated that the coefficient for hexadirectional orientation stability was larger than that for hexadirectional spatial consistency (Δβ = 0.369, 95% CrI [0.112, 0.628]). Although the coefficient for hexadirectional orientation stability was also numerically larger than that for temporal hexadirectional geometry stability, this difference remained uncertain (Δβ = 0.147, 95% CrI [−0.152, 0.446]). Thus, hexadirectional orientation stability was the most consistently supported predictor of pmEC hexadirectional signal across hemispheres, although its distinction from hexadirectional geometry stability was less clear in right-pmEC.

Because hexadirectional signal and hexadirectional orientation stability both incorporate cross-run comparisons of matching direction bins, their association could partly reflect shared computation. We therefore conducted a control analysis in which hexadirectional signal was recalculated after excluding these shared comparisons, and the joint model was repeated (See Method). In left-pmEC, hexadirectional orientation stability remained the only stability metric clearly associated with hexadirectional signal, (β = 0.267, 95% CrI [0.054, 0.475]). In right-pmEC, hexadirectional orientation stability was not clearly associated with the off-diagonal hexadirectional measure. The remaining predictors showed the same overall pattern as in the primary analysis (Table S1).

Finally, we tested whether hexadirectional orientation stability was behaviorally relevant for object-location memory. In an age-adjusted binomial model, higher left-pmEC hexadirectional orientation stability was associated with better object-location accuracy (Figure 2E; log-odds coefficient= 0.158, 95% CrI [0.018, 0.299]). Chronological age was negatively associated with accuracy (log-odds coefficient = - 0.295, 95% CrI [−0.439, −0.148]), indicating that the behavioral measure was sensitive to age-related differences. In right-pmEC, hexadirectional orientation stability showed no clearly supported association with object-location accuracy (log-odds coefficient = −0.025, 95% CrI [−0.168, 0.113]), although age again showed a negative association with accuracy (log-odds coefficient =-0.295, 95% CrI [-0.429, −0.148]). Neither left-nor right-pmEC hexadirectional signal showed a clear association with accuracy (Table S2).

Together, the primary analyses identified hexadirectional orientation stability as a predictor of pmEC hexadirectional signal in both hemispheres. However, in the control analysis, robust evidence for this stability-signal association was observed only in left-pmEC, and hexadirectional orientation stability was also associated with object-location accuracy. These findings identified left-pmEC as the most robust and behaviorally relevant locus of the effect, motivating the subsequent circuit-level analyses.**Figure 2**. (A)

### Local pmEC structural properties do not explain hexadirectional signal strengths or orientation stability

We next asked whether individual differences in left-pmEC hexadirectional signal and hexadirectional orientation stability were associated with local structural properties: mean diffusivity, a marker of gray-matter microstructural integrity; T1w/T2w ratio, a myelin-sensitive proxy; and pmEC volume (controlled for total intracranial volume). In age-adjusted models, hexadirectional signal showed no clear association with mean diffusivity (β = −0.105, 95% CrI [−0.356, 0.148], BF₀₁ = 2.77), T1w/T2w ratio (β = −0.143, 95% CrI [−0.366, 0.079], BF₀₁ = 1.96), or pmEC volume (β = 0.178, 95% CrI [−0.035, 0.390], BF₀₁ = 1.16). Hexadirectional orientation stability likewise showed no clear association with mean diffusivity (β = −0.013, 95% CrI [−0.274, 0.243], BF₀₁ = 3.85), T1w/T2w ratio (β = −0.039, 95% CrI [−0.262, 0.185], BF₀₁ = 4.20), or pmEC volume (β = 0.069, 95% CrI [−0.142, 0.279], BF₀₁ = 3.79). For hexadirectional orientation stability, all Bayes factors provided moderate evidence against an association (BF₀₁ > 3), making it less likely that its variability simply reflects the measured local structural differences in pmEC. The same overall pattern was observed in right-pmEC (Table S3).

### Retrosplenial orientation stability is related to hexadirectional orientation stability and hexadirectional signal magnitude in left pmEC

Having found that neither age nor local pmEC structural properties clearly explained either left-pmEC measure, we next asked whether hexadirectional orientation stability was related to the stability of orientation signals in retrosplenial cortex and presubiculum (Figure 3A), two regions known to provide directional inputs to entorhinal cortex^18,21–24^. Retrosplenial absolute-orientation stability (see Methods) was associated with left-pmEC hexadirectional orientation stability, (Figure 3B; β = 0.221, 95% CrI [0.012, 0.423]). The corresponding association with presubicular absolute-orientation stability was weaker and more uncertain, (Figure 3C; β = 0.175, 95% CrI [−0.041, 0.383]). Moreover, retrosplenial absolute-orientation stability was also associated with object-location accuracy (Figure 3D; log-odds = 0.224, 95% CrI [0.080, 0.368]). Together, these findings indicate that retrosplenial absolute-orientation stability is linked to both left-pmEC hexadirectional orientation stability and object-location accuracy, consistent with a circuit-level account in which reliable orientation-related signals may support the temporal stability of entorhinal hexadirectional representations in older adults.

**Figure 3.**
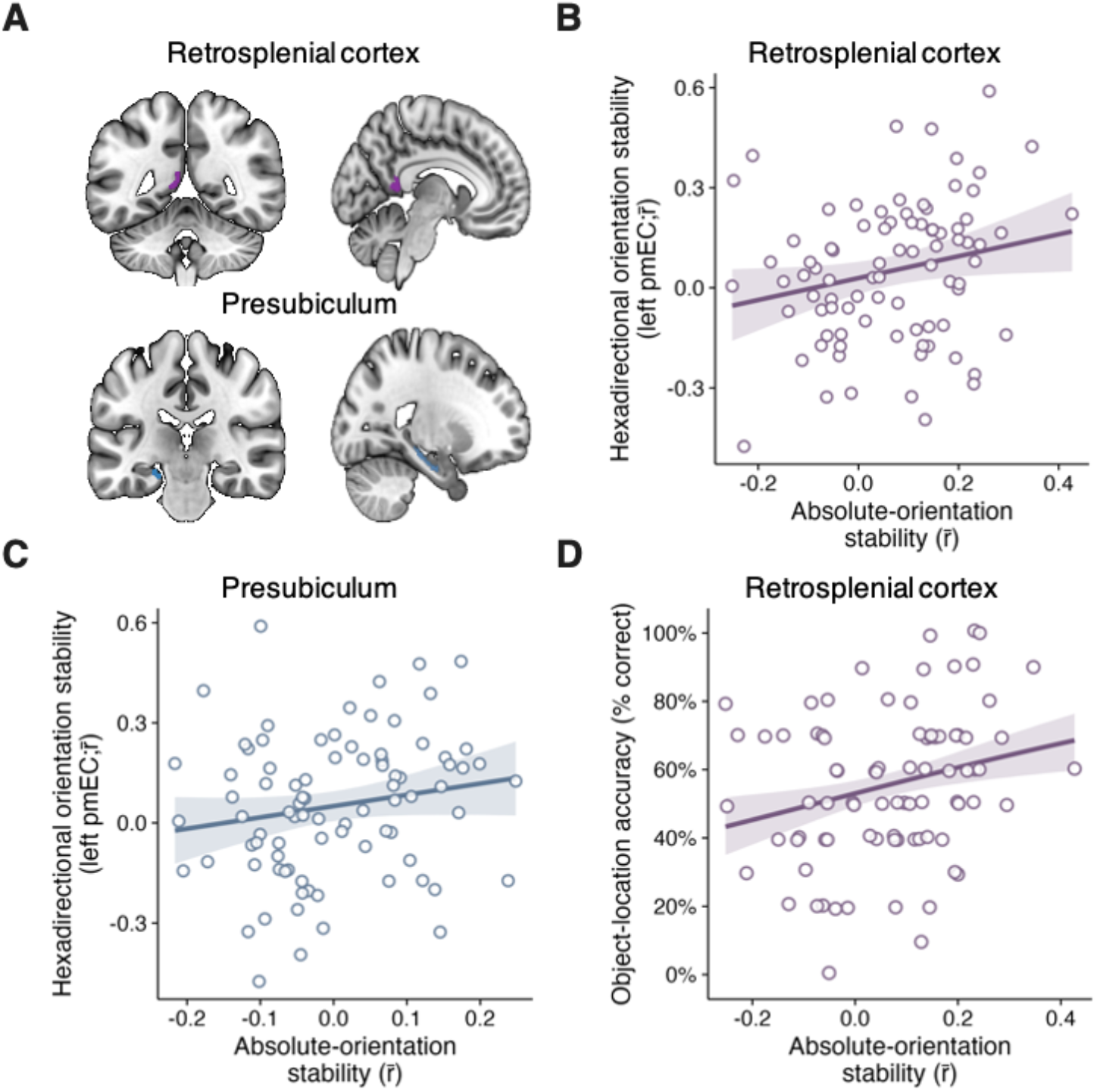
(A) Left retrosplenial cortex (purple) and presubiculum (blue) regions of interest. (B,C) Associations between absolute-orientation stability in retrosplenial cortex (B) or presubiculum (C) and left-pmEC hexadirectional orientation stability. (D) Retrosplenial absolute-orientation stability was associated with object-location accuracy in an age-adjusted binomial model. Lines show fitted age-adjusted linear relationships.

Finally, we tested a more specific prediction of this account. If retrosplenial absolute-orientation stability contributes to hexadirectional representations by stabilizing their temporal expression in left-pmEC, then left-pmEC hexadirectional orientation stability should statistically link retrosplenial absolute-orientation stability to left-pmEC hexadirectional signal. Mediation analysis was consistent with an indirect association between retrosplenial absolute-orientation stability and hexadirectional signal through hexadirectional orientation stability (Figure 4; indirect effect = 0.125, 95% CrI [0.008, 0.261], Table S4).

**Figure 4.**
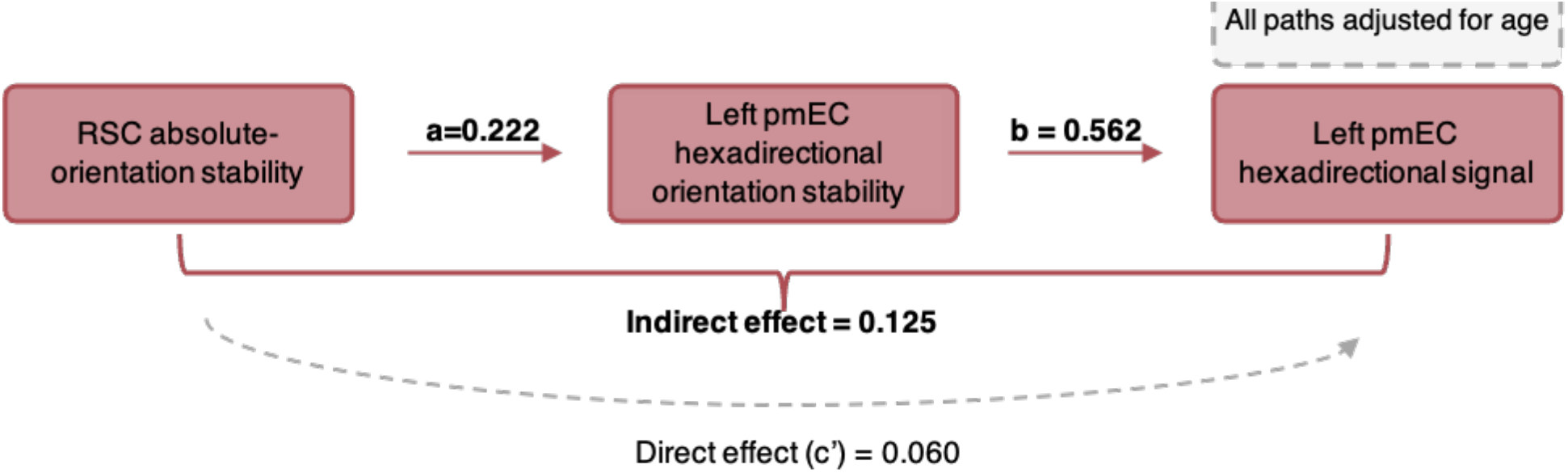
Age-adjusted Bayesian mediation model. The *a* and *b* paths denote the associations of retrosplenial absolute-orientation stability with left-pmEC hexadirectional orientation stability and of left-pmEC orientation stability with hexadirectional signal, respectively. The indirect effect estimates the association statistically accounted for by left-pmEC orientation stability; the dashed path denotes the direct association. Statistically reliable effects are in **bold**.

## DISCUSSION

In this study, we investigated candidate neural correlates of individual differences in entorhinal hexadirectional coding in typical aging by combining virtual-navigation fMRI and multivariate representational similarity analysis with structural and diffusion MRI and an object-location-memory assessment. Hexadirectional orientation stability predicted hexadirectional signal strength in both pmEC hemispheres, whereas chronological age, the local structural measures examined, hexadirectional geometry stability, and hexadirectional spatial consistency showed no clear associations. Evidence converged in left pmEC: in a control analysis excluding shared matching-bin comparisons, the association between hexadirectional signal and orientation stability persisted, and higher orientation stability was also associated with better object-location memory, whereas hexadirectional signal magnitude itself showed no clear association with memory. Furthermore, retrosplenial absolute-orientation stability was associated with both left-pmEC hexadirectional orientation stability and memory performance. Mediation analysis supported an indirect association between retrosplenial absolute-orientation stability and left-pmEC hexadirectional signal through left-pmEC hexadirectional orientation stability. Together, these findings suggest that individual differences in entorhinal hexadirectional coding in aging may partly reflect how reliably grid-relative orientation patterns are maintained over time, potentially linked to the stability of retrosplenial orientation representations.

The marked variability in pmEC hexadirectional signal is consistent with the broader heterogeneity of cognitive and neural functioning in later life^25–28^. Within spatial navigation, Stangl et al.^7^ found that some older adults retained hexadirectional signals and path-integration performance comparable to those of younger adults despite lower group-average values. A similar pattern of reduced average spatial learning together with greater inter-individual variability has been reported in rats^29^. Thus, group-level age effects can coexist with marked heterogeneity in spatial function; age alone does not explain why grid-like coding is preserved in some older adults but weakened in others. Moreover, hexadirectional signal was not associated with local pmEC micro-(structure), including volume, myelin-sensitive T1w/T2w contrast, or mean diffusivity. Although these MRI measures may not capture subtler cellular, synaptic, or layer-specific alterations, the findings direct attention to the functional organisation of the representation itself.

To identify the type of representational stability most relevant to grid-like coding, we compared three complementary properties of the stability of the pmEC signal. Of the three properties examined, hexadirectional orientation stability was the only one consistently associated with hexadirectional signal across hemispheres. Furthermore, in left-pmEC its association was larger than those for hexadirectional geometry stability and spatial consistency. This specificity distinguishes between forms of representational instability: hexadirectional geometry stability can remain preserved even if the mapping between environmental directions and grid-relative axes shifts across runs provided that the similarity structure among direction-specific patterns is maintained, whereas hexadirectional spatial consistency indexes the dispersion of response amplitudes across voxels rather than their alignment to common grid axes. Importantly, higher left-pmEC hexadirectional orientation stability was associated with better object-location memory, whereas hexadirectional-signal magnitude showed no clear association with accuracy. Because participants had to identify learned locations across different viewpoints, stable grid-relative orientation patterns may provide the common spatial reference required for retrieval; if these patterns drift, a location may instead be reinstated relative to a shifted reference frame.

This joint neural and behavioral specificity substantially refines our understanding of temporal instability as observed in previous work^7^. Given that directions 60° apart were assigned to the same grid-relative bin, a drift in their alignment across runs would reduce both pattern reproducibility and fit to the hexadirectional model without necessarily erasing the underlying spatial organization. Relative grid-cell phases can remain stable despite shifts in the absolute phases of individual cells^30^. Similarly, aged mouse medial entorhinal cortex retains identifiable grid cells and coordinated spatial maps, but firing stability is impaired and, unlike grid-cell density, is related to spatial-memory performance^10^. Together, these findings support a mechanism by which aging weakens the reliable environmental alignment and behavioral usefulness of organized spatial representations, rather than simply eliminating them.

One possible source of instability in the pmEC grid-relative orientation code is variability in the incoming orientation signals used to anchor it to the environment. We therefore assessed the cross-run reproducibility of absolute orientation-related patterns in retrosplenial cortex and presubiculum in full 360° space and found that retrosplenial absolute-orientation stability was associated with left-pmEC hexadirectional orientation stability and object-location memory. Critically, mediation analysis supported an indirect association between retrosplenial absolute-orientation stability and left-pmEC hexadirectional signal through left-pmEC hexadirectional orientation stability. Retrosplenial cortex is well positioned to provide a stable environmental orientation reference because it integrates landmark and scene information with internally generated directional signals. In mice, RSC neurons encode orientation relative to the environment without physical head movement and combine visual and thalamic inputs in a manner consistent with landmark-based heading anchoring^31^. In humans, retrosplenial fMRI activity patterns encode heading within both global and local environmental reference frames^18,32^. Against this background, our findings suggest that the reliable reinstatement of orientation-specific RSC patterns may be relevant to the temporal stability of pmEC grid-relative orientation patterns, as well as hexadirectional signal and object-location memory in older adults.

The absence of a similarly robust association with presubicular absolute-orientation stability may seem surprising because retrosplenial orientation-related information is often thought to be integrated within presubicular directional circuits before reaching entorhinal cortex^33,34^. However, it may reflect both anatomy and measurement. In primates, RSC also shares direct projections with entorhinal cortex^35^, and suppressing retrosplenial afferents in layer 3 of dorsal medial entorhinal cortex in mice reduces visually evoked responses^36^, indicating that orientation-related input need not be exclusively relayed through the presubiculum. In addition, the substantially smaller presubicular ROI is more susceptible to partial-volume effects and segmentation or registration inaccuracies, which could reduce the reliability of its multivariate stability estimate. Accordingly, the present data do not resolve the anatomical pathway through which orientation-related information relates to pmEC hexadirectional orientation stability but identify RSC as the region in which this relationship was most clearly evident.

What mechanisms could modulate the stability of orientation signals in retrosplenial cortex? Recent behavioral and computational work provides evidence that older adults make larger angular errors and accumulate greater heading uncertainty over short delays, with increased synaptic noise identified as one mechanism capable of reproducing these effects in a ring-attractor model^19^. Direct recordings of head-direction coding in normally aged rodents remain limited. Nonetheless, aged mice showed diminished innervation and synaptic drive in the medial-vestibular-nucleus–dorsal-tegmental-nucleus pathway, and activating this pathway improved navigation^37^, indicating that head-direction circuitry is vulnerable in normal aging. In a mouse model of pathological tau, direct recordings further showed reduced directionality and directional coherence of head-direction cells^38^. Attractor dynamics can preserve an organized orientation code while allowing deviations along the heading manifold to persist unless external information corrects them^39^. Visual landmarks and environmental boundaries provide such correction for head-direction^40^ and grid-cell activity^41^, respectively. Accordingly, inconsistent RSC orientation-related patterns could provide a less reliable corrective reference, allowing the mapping between environmental directions and grid-relative orientations to drift across runs. Directions that should be equivalent in hexadirectional space may then become offset, reducing pmEC hexadirectional orientation stability and attenuating the aggregate hexadirectional signal.

Several limitations apply to these conclusions. Navigation was visually simulated and therefore lacked the vestibular and proprioceptive information available during natural movement, which may have increased orientation uncertainty and magnified inter-individual differences in hexadirectional orientation stability. During natural navigation, vestibular and proprioceptive cues may help constrain drift, particularly in aging when individual sensory signals become noisier^42,43^. Future work using multisensory navigation paradigms will be needed to determine whether retrosplenial orientation-related stability contributes less to pmEC stability and hexadirectional signal under these conditions. Because the sample included only older adults, it also remains unclear whether these associations are amplified in aging or reflect individual differences present across adulthood. Younger-adult comparisons and longitudinal multisensory-navigation studies will be needed to address this question. Finally, the object-location task comprised relatively few trials and therefore provides a coarse functional readout rather than a detailed assessment of the behavioral consequences of orientation instability. Future work should combine more targeted orientation-maintenance tasks with trial-wise estimates of angular drift to test how neural orientation stability relates to spatial memory deficits in aging.

These findings have implications for how age-related reductions in hexadirectional coding are conceptualized and potentially targeted. Rather than reflecting the disappearance of an underlying grid code, weaker hexadirectional signal may arise when that representation cannot be maintained reliably enough over time to be consistently expressed at the population level. Such instability could affect memory as well as navigation, because grid cells have been proposed to help anchor experiences in space^44^ and organize relationships within episodic memories^45^. Unstable representations could therefore compromise the binding of items and their contextual relationships, reducing the coherence of encoding and subsequent reinstatement. Importantly, this account reframes age-related variation in grid-like coding as a potentially modifiable problem of representational reliability: a preserved representation may still be expressed reliably when environmental landmarks and convergent self-motion cues provide a sufficiently stable reference. Furthermore, our retrosplenial findings point to the stability of orientation inputs as one potentially important contributor to this temporal reliability. Relevant computations appear modifiable in animal work: enhancing GABAergic signaling partially restored visual cortical orientation and direction selectivity in senescent macaques^46^, whereas activation of a vestibular pathway within the head-direction network ameliorated navigation deficits in aged mice^37^. Although these interventions do not directly establish that entorhinal coding can be stabilized, they motivate testing whether strengthening landmark-based orientation processing or better integration of visual and self-motion cues can improve the stability of grid-like representations and support navigation and memory in older adults.

Together, these findings move beyond documenting weaker grid-like coding in older age to identify a specific form of representational instability and a candidate circuit-level contributor: the reliability of retrosplenial orientation-related representations. The convergence of pmEC stability, retrosplenial absolute-orientation stability, and object-location-memory findings points to a functional organization in which stable orientation-related representations may help maintain a common spatial reference across experience. In this account, age-related variation in grid-like coding is not simply a matter of whether a spatial representation is present, but of whether it can be reliably reinstated when needed. Future work combining direct measures of heading maintenance with neural measures of representational stability will be important for testing this account more directly.

## RESOURCE AVAILABILITY

### Lead contact

Requests for further information and resources should be directed to and will be fulfilled by the lead contact, Vladislava Segen.

### Materials availability

This study did not generate new unique reagents.

### Data and code availability

- Data reported in this paper will be shared by the lead contact upon request.
- Original analysis code for generation of hexadirectional signal and stability metrics is available at 10.5281/zenodo.23106522
- Any additional information required to reanalyze the data reported in this paper is available from the lead contact upon request.

## ACKNOWLEDGMENTS

This work was supported by the VELUX Stiftung (Project No. 1809) and the German Research Foundation (Deutsche Forschungsgemeinschaft, DFG) under Project ID 425899996 (CRC 1436) and by BrightFocus foundation (A2026010F), which supported V.S. We also gratefully acknowledge the research assistants who contributed to project administration and data collection.

## AUTHOR CONTRIBUTIONS

VS: Conceptualization, Methodology, Software, Formal analysis, Data curation, Visualization, Project administration, Funding acquisition, Writing – original draft, Writing – review & editing. GCN: Formal analysis, Writing – review & editing. PB: Formal analysis, Project administration, Writing – review & editing. TBB: Conceptualization, Methodology, Project administration, Writing – review & editing. JIS: Investigation, Writing – review & editing. TW: Conceptualization, Funding acquisition, Project administration, Supervision, Writing – review & editing.

## STAR★METHODS

### Key resources table

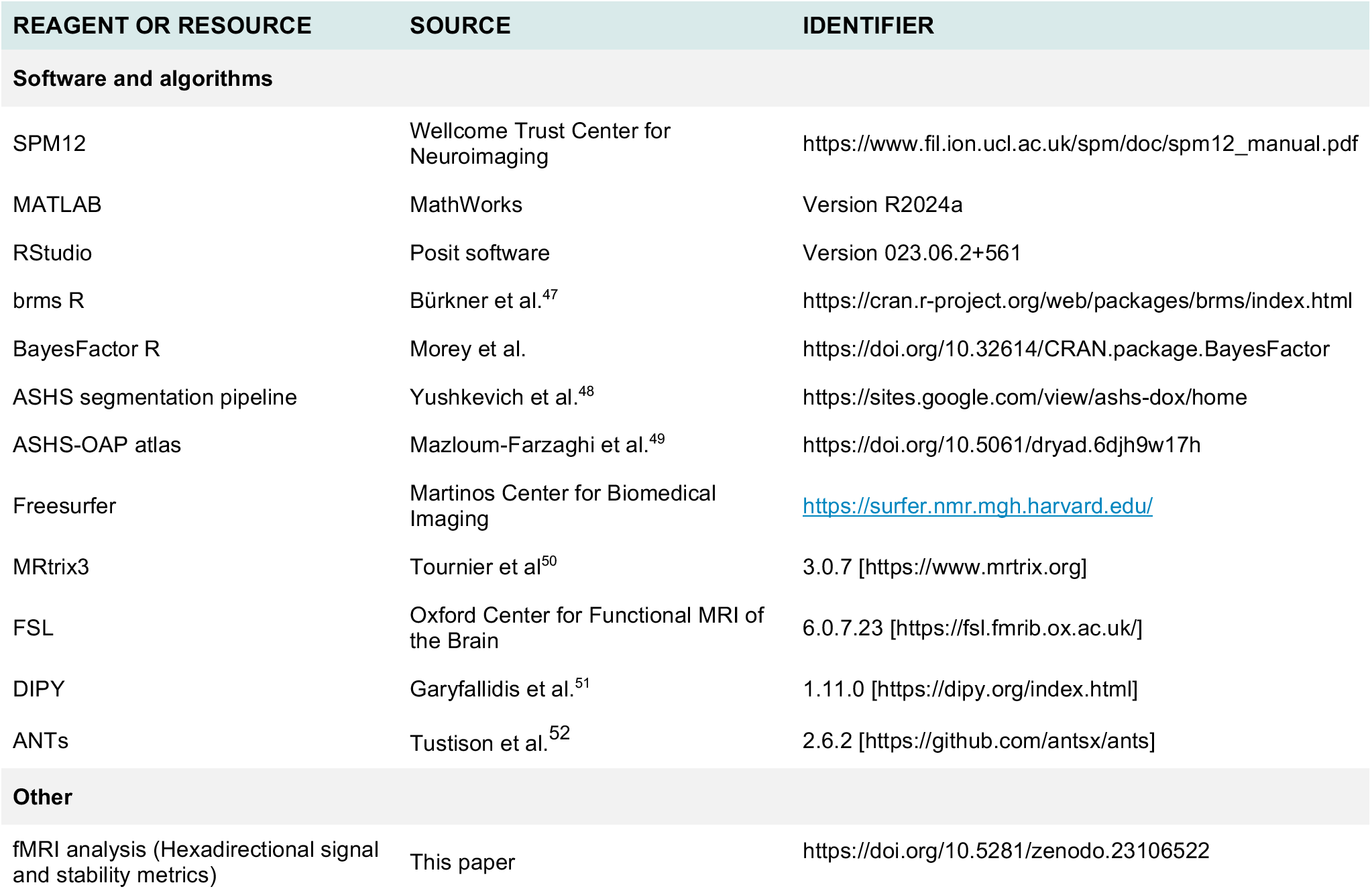

### 1. Experimental model and study participant details

#### Human participants

The present investigation was conducted as a subproject within a larger cohort study of healthy aging^52^. Participants were recruited primarily from the main healthy-aging cohort, supplemented by recruitment through related studies within the broader research programme. The sample used in this study consisted of 88 cognitively unimpaired, community-dwelling older adults. Cognitive status was assessed using the Consortium to Establish a Registry for Alzheimer’s Disease Neuropsychological Battery Plus (CERAD-Plus)^53^, where available. Participants were excluded if they scored below 26 on the Mini-Mental State Examination or if their global CERAD-Plus performance fell below −1.5 SD relative to age-, sex-, and education-adjusted norms (mean=0.39, sd=0.49). This resulted in removal of 1 participant (MMSE below 26). Five participants did not complete the CERAD-Plus battery but completed the Montreal Cognitive Assessment^54^ and scored above 23, consistent with normal cognition^55^.

Additionally, one participant was excluded because the entorhinal cortex could not be reliably segmented due to atypical anatomy, and one participant was excluded because visual problems affected task performance. The final fMRI sample comprised 85 participants (41 female; mean age = 75.39 years, SD = 7.93, range = 62-96 years). Diffusion weighted MRI data was only available for a subset of participants (61; 30 female, mean age = 74.39 years, SD = 7.69, range=62-87).

### 2. Method details

#### fMRI grid-cell task

Participants performed a passive virtual navigation task designed to sample a broad range of travel directions for the assessment of hexadirectional representations. In each run, participants were translated along one of nine predefined trajectory versions, which were pseudorandomized across participants. Each trajectory comprised translation periods, rotations, and brief stationary intervals, producing a random-walk-like path through the environment. Translation speed was held constant at 15 virtual m/s, rotations were performed at 20°/s, and translation segments varied in duration (mean = 3.45 s, SD = 0.75). Depending on the assigned trajectory, each run lasted between 8.5 and 9.5 min.

The virtual environment consisted of a square room with textured flooring and wall cues providing orientation information. Five everyday objects were distributed across the environment and marked by red cones to facilitate visibility. In the second run, the spatial layout remained identical, but with new target objects and locations. Participants were instructed to imagine actively moving through the environment and to encode the object locations. After each run, they completed 5 trials of a three-alternative forced-choice object-location test, selecting the screenshot that showed the correct location of each target object, in total resulting in 10 trials object location memory per participant.

#### MRI acquisition

Structural MRI and fMRI data were acquired on a 3T Siemens Prisma scanner using a 64-channel head coil. A whole-brain T1-weighted MPRAGE image was acquired with TR = 2,500 ms, TE = 4.37 ms, TI = 1,100 ms, flip angle = 7°, and 1.0mm isotropic resolution. A coronal T2-weighted turbo spin echo scan, oriented perpendicular to the hippocampal long axis, was acquired for entorhinal and medial temporal lobe segmentation (TR = 6,000 ms, TE = 71 ms, 0.5 × 0.5 × 2.0 mm resolution).

Functional images were acquired using a two-echo 3D EPI sequence with TR = 2,200 ms, TE1 = 14.2 ms, TE2 = 39.2 ms, flip angle = 15°, 2.0 mm isotropic resolution, 80 slices, and CAIPIRINHA acceleration of 3 × 2. Two task runs were acquired, each comprising 276 volumes.

Diffusion MRI data, available for a subset of participants (n=61), were acquired in a separate session on a Siemens Skyra scanner using a 32-channel head coil. Diffusion-weighted images were acquired using a single-shot echo-planar imaging sequence with TR = 4,010 ms, TE = 72.4 ms, 2.0 mm isotropic resolution, field of view = 240 × 240 × 144 mm3, two-fold parallel acceleration, and partial Fourier factor = 7/8. The acquisition included 70 volumes with b-values of 0, 700, and 1,000 s/mm2, comprising 10 b = 0 volumes and 30 diffusion directions for each non-zero b-value.

#### Region of interest masks

The posterior-medial entorhinal cortex (pmEC) was segmented using the Automatic Segmentation of Hippocampal Subfields (ASHS)^57^ pipeline with the ASHS-OAP atlas^49^, an atlas developed from older adults for automated segmentation of MTL cortical regions and hippocampal subfields. The pipeline used participants’ high-resolution T2-weighted images together with the corresponding T1-weighted images and generated participant-specific masks of pmEC in each hemisphere. An experienced rater visually quality-controlled the resulting masks on the high-resolution T2-weighted images according to the manual segmentation protocol of Olsen et al^58^. All masks passed visual quality control (after exclusion of the participant with the abnormal anatomy). The pmEC was selected a priori because it is considered the putative human homologue of the rodent medial entorhinal cortex, where grid cells were first identified and have been most extensively characterized^5,59^. Human fMRI studies of hexadirectional coding have also commonly focused on this entorhinal subdivision^60–62^. Hexadirectional signal, hexadirectional stability metrics, and structural control measures were extracted separately from participant-specific left and right pmEC masks in native space.

The left presubiculum and retrosplenial cortex were included as candidate orientation-related input regions since both are implicated in orientation, heading, and spatial reference-frame processing^18,21–24^ and could therefore provide signals that help maintain the alignment of entorhinal hexadirectional representations over time^14^. The left presubiculum was manually delineated on the high-resolution T2-weighted images following the anatomical guidelines described by Dalton et al^63^. All presubiculum masks were independently cross-checked by a second rater. Retrosplenial cortex was approximated using the left ventral posterior cingulate cortex region (vPCC) from the Destrieux atlas^64^ implemented in FreeSurfer. Participant-specific vPCC masks were obtained from the cortical parcellation derived from the T1-weighted anatomical images and were used as an anatomical proxy for retrosplenial cortex.

#### fMRI preprocessing

Functional MRI preprocessing was performed using SPM12^65^ in MATLAB. Motion parameters were estimated from the first echo across runs using a two-pass register-to-mean procedure. The resulting transformations were applied to the corresponding second-echo volumes, and both echoes were subsequently resliced to a common space defined by the first retained volume of the first run. The two echoes were combined using voxel-wise weights derived from a brief resting-state scan (80 volumes) acquired with the same sequence parameters. For each echo, temporal signal-to-noise ratio maps were computed and used to generate voxel-wise echo-combination weights. These weights were applied to each task volume, yielding one optimally combined functional time series per run. The combined images were spatially smoothed with a 5 mm full-width-at-half-maximum Gaussian kernel.

Physiological noise regressors were derived using an anatomical CompCor approach^66^. The T1-weighted image was coregistered to the mean EPI image and segmented into gray matter, white matter, and cerebrospinal fluid using SPM’s unified segmentation procedure. The first five principal components extracted from white matter and cerebrospinal fluid masks were retained as nuisance regressors.

To align the anatomical regions of interest with the functional data, registration was performed separately according to the anatomical image from which each ROI was derived. For the T2-derived pmEC and presubiculum masks, the high-resolution T2-weighted image was coregistered directly to the mean EPI image, and the resulting transformation was applied to the corresponding masks to bring them into native functional space. For the T1-derived vPCC masks used to approximate retrosplenial cortex, the transformation estimated during coregistration of the T1-weighted image to the mean EPI image was applied to bring the masks into native functional space. All masks were resampled using nearest-neighbour interpolation to preserve their discrete anatomical labels and were visually inspected for alignment with the functional images.

### First-level modelling for hexadirectional signal and stability analyses

First-level general linear models included direction-specific regressors that were defined from periods of forward translation through the virtual environment. For each translation segment, the instantaneous movement direction was expressed as an angular heading and folded into 60° rotational space, yielding six direction regressors corresponding to 0°, 10°, 20°, 30°, 40°, and 50°. Regressors were modelled using the onset and duration of the corresponding translation periods and convolved with the canonical hemodynamic response function. Rotations and stationary intervals were not used to estimate direction-specific beta patterns. Each model additionally included the six realignment parameters and five anatomical CompCor components as nuisance regressors. Model estimation yielded one beta image for each direction regressor in each run.

#### Multivariate hexadirectional signal in pmEC

Hexadirecitonal signal was assessed using multivariate representational similarity analysis applied to the direction-specific beta images generated by the preceding first-level GLM. For each participant and pmEC ROI, voxel-wise beta estimates for the six direction regressors were extracted separately from run 1 and run 2. We then computed a cross-run neural representational dissimilarity matrix by comparing each direction-specific multivoxel pattern from run 1 with each direction-specific pattern from run 2 using 1-Pearson correlation. This yielded a 6 × 6 neural RDM for each ROI, in which each cell reflected the dissimilarity between two movement-direction bins estimated from independent runs.

To test whether this neural representational structure followed the hexadirectional symmetry expected for hexadirectional coding, we constructed a model RDM based on the angular difference between movement-direction bins. Angular differences were transformed using a cosine function with hexadirectional periodicity, such that directions separated by multiples of 60° were predicted to have more similar activity patterns than pairs separated by angular offsets that were not multiples of 60°: RDMmodel = 1 - cos(6Δθ). The neural and model RDMs were vectorized and compared using Spearman rank correlation. The resulting correlation coefficient served as the participant-specific estimate of hexadirectional signal strength for left and right pmEC.

### Control analysis

The association between hexadirectional signal and hexadirectional orientation stability could be partly inflated because both measures incorporate cross-run comparisons between matching direction bins. Hexadirectional orientation stability quantifies the similarity of grid-relative orientation patterns across runs, and these same six matching-bin pairs form the diagonal elements of the cross-run neural RDM used to estimate hexadirectional signal. To control for this potential computational overlap, we recalculated hexadirectional signal after excluding these six diagonal elements from both the 6 × 6 neural RDM and the corresponding model RDM before vectorization, and calculated the Spearman correlation using the remaining 30 cross-run comparisons.

#### Temporal and spatial stability of pmEC hexadirectional signal

The direction-specific beta images generated by the same first-level GLM were also used to derive complementary measures of the temporal and spatial stability of pmEC hexadirectional coding.

*Hexadirectional geometry stability* was assessed by computing separate run-specific neural RDMs based on pairwise correlation distances between the six direction-specific beta patterns within each run. The off-diagonal elements of the run 1 and run 2 RDMs were vectorized and correlated across runs using Spearman correlations. This model-free measure captured the temporal reliability of direction-related representational geometry independently of correspondence with the six-fold model RDM.

*Hexadirectional orientation stability* was defined as the run-to-run consistency of direction-specific multivoxel activity patterns in 60° rotational space. For each of the six direction bins, we compared the corresponding direction-specific beta pattern from run 1 with the matching beta pattern from run 2 within the pmEC ROI. The Pearson correlations were then z-transformed and averaged across the six bins to provide an index of how consistently the six-fold orientation-specific pattern was expressed across runs.

*Hexadirectional spatial consistency* was assessed from the spatial distribution of direction-specific beta estimates within pmEC. For each 60° direction bin, we quantified within-ROI beta standard deviation across voxels and averaged this measure across bins and runs. Higher within-ROI beta standard deviation indicated lower spatial stability of the direction-specific pattern.

#### Absolute orientation stability in left RSC and presubiculum

To quantify the temporal stability of absolute orientation coding in the left retrosplenial cortex and left presubiculum, we estimated a separate first-level GLM in which movement directions were represented across the full 360° space rather than folded into 60° rotational space, as we did not expect hexadirectional signal in these regions. Forward-translation periods were assigned to 12 orientation bins centered at 30° intervals (0°, 30°, 60°,…, 330°). Each bin included movement directions within ±10° of its central orientation, with circular wrapping at 0°; for example, the 0° bin comprised directions of 350°, 0°, and 10°. All other model specifications were identical to those of the hexadirectional-related first-level GLM.

*Absolute-orientation stability* was quantified separately within each candidate region by correlating the multivoxel beta pattern for each of the 12 orientation bins between runs 1 and 2 using Pearson correlation. The resulting 12 correlations were averaged to obtain one orientation-stability estimate per participant and region. This measure indexed the extent to which absolute heading-specific activity patterns were preserved across runs, independently of the six-fold rotational structure used to quantify hexadirectional representations.

#### Structural and diffusion MRI preprocessing and analysis

To determine whether individual differences in pmEC hexadirectional signal hexadirectional orientation stability and were related to local anatomical properties, we derived three complementary structural measures: gray-matter volume, the T1w/T2w ratio, and diffusion-derived mean diffusivity (MD). Each measure was extracted separately from the participant-specific left and right pmEC ROIs.

Gray-matter-constrained pmEC volumes were obtained from the ASHS segmentations generated using the `UseGray` option. Because regional brain volumes covary with overall head size, raw pmEC volume was adjusted for total intracranial volume (TIV). Separately for each hemisphere, raw pmEC volume was regressed on TIV across participants, and the resulting residuals were retained. These residualised estimates reflected variation in pmEC volume beyond that attributable to individual differences in head size and were used in the subsequent statistical analyses.

The T1w/T2w ratio was used as an MRI-based proxy for intracortical myelination, which is commonly interpreted as being sensitive to intracortical myelin content^63^. The T1-weighted image was processed using SPM12 unified segmentation, which generated gray-matter, white-matter, and cerebrospinal-fluid tissue classes together with a bias-field-corrected T1-weighted image. The corresponding T2-weighted image was rigidly coregistered to the T1-weighted image using normalized mutual information, resliced using fourth-order B-spline interpolation, and corrected for intensity nonuniformity. Before calculating the ratio, the intensity of each bias-corrected image was normalized by the median signal within its corresponding white-matter mask. The normalized T1-weighted image was then divided voxelwise by the normalized T2-weighted image, with calculation restricted to gray matter. The pmEC masks were transferred to the processed T1-weighted image using nearest-neighbour interpolation, and the mean T1w/T2w ratio was extracted from each ROI. Higher values were interpreted as indicating greater relative intracortical myelin-sensitive contrast.

Diffusion-weighted MRI data were preprocessed using tools from MRtrix3 and FSL. Processing included thermal-noise reduction, correction of susceptibility-induced geometric distortions using b0 images acquired with opposing phase-encoding directions, and correction of participant motion and eddy-current-induced distortions. Bias-field correction was performed using the ANTs-based algorithm implemented in MRtrix3’s `dwibiascorrect`, and brain masks were generated using FSL’s Brain Extraction Tool. Data quality following eddy-current and motion correction was evaluated using `eddy_quad`.

Diffusion tensors were fitted to the preprocessed data using `dipy.reconst.dti.TensorModel` from DIPY, producing participant-specific MD maps. To transfer the anatomical ROIs into diffusion space, the high-resolution T2-weighted image was registered to the participant’s mean b0 image, and the resulting transformation was applied to the pmEC masks before resampling them onto the diffusion grid. Mean pmEC MD was calculated using the resampled mask values as partial-volume weights, including all voxels withhex a nonzero mask contribution.

### 3. Quantification and statistical analysis

All statistical analyses were conducted in RStudio. Bayesian linear regression models were fitted using the *brms*^43^ package with a Gaussian likelihood. Continuous predictors, outcomes, and covariates were z-standardized before model estimation unless otherwise stated, and age was included as a covariate where appropriate. Results from Gaussian models were reported as standardized beta coefficients with 95% credible intervals. The primary model tested whether pmEC hexadirectional signal was predicted by hexadirectional orientation, geometry and spatial stability stability. Additional models tested whether pmEC structural measures—volume, T1w/T2w ratio, and mean diffusivity—explained pmEC hexadirectional signal beyond age. To examine whether the stability of absolute-orientation inputs was associated with pmEC hexadirectional orientation stability, separate regression models tested the relationships of RSC and presubiculum absolute-orientation stability with pmEC hexadirectional orientation stability. Where applicable, Bayes factors were calculated by comparing each full model against an age-only or intercept-only reference model.

Behavioural relevance was assessed using Bayesian binomial regression models with a logit link. The number of correct responses on the three-alternative forced-choice object-location task was modelled relative to the number of trials answered across both runs. Separate age-adjusted models tested whether pmEC hexadirectional orientation stability, pmEC hexadirectional-signal magnitude, and RSC absolute-orientation stability predicted object-location performance, with analyses conducted separately by hemisphere where applicable. Continuous predictors and age were z-standardized, whereas the behavioural outcome was retained as a binomial count. Results from these models were reported as log-odds coefficients with 95% credible intervals.

A Bayesian mediation analysis was used to examine whether the association between RSC absolute-orientation stability and left pmEC hexadirectional signal was statistically mediated by left pmEC hexadirectional orientation stability. Two Bayesian Gaussian regression models were fitted: a mediator model predicting pmEC hexadirectional orientation stability from RSC absolute-orientation stability and age, and an outcome model predicting pmEC hexadirectional signal from pmEC hexadirectional orientation stability, RSC absolute-orientation stability, and age. Weakly informative priors were used for all regression coefficients, β ∼ Normal(0, 1), with Student-t(3, 0, 2.5) priors for intercepts and Exponential(1) priors for residual standard deviations. Models were fitted with four chains, 4,000 iterations per chain, and 1,000 warm-up iterations, using an adapt_delta value of 0.95. The coefficient relating RSC absolute-orientation stability to pmEC hexadirectional orientation stability represented the **a** path, whereas the coefficient relating pmEC hexadirectional orientation stability to hexadirectional signal, conditional on RSC absolute-orientation stability, represented the **b** path. The indirect effect was computed for each posterior draw as the product of the corresponding **a** and **b** path samples. The direct effect (**c’**) represented the association between RSC absolute-orientation stability and pmEC hexadirectional signal after accounting for the mediator, and the total effect was calculated as (**c’ + ab**). The posterior distributions of the direct, indirect, and total effects were summarized using the posterior median, 95% credible interval, and probability of direction.

#### Declaration of generative AI and AI-assisted technologies in the manuscript preparation process

During preparation of this work, the authors used ChatGPT (OpenAI) for language editing and to assist with the development and troubleshooting of analysis code. All analyses, code validation, interpretation, and final decisions were performed and verified by the authors. The author(s) reviewed and edited the output as needed and take full responsibility for the content of the published article.

## REFERENCES

1. Moffat, S.D., and Resnick, S.M. (2002). Effects of age on virtual environment place navigation and allocentric cognitive mapping. Behavioral Neuroscience 116, 851–859. 10.1037/0735-7044.116.5.851.

2. Zhou, R., Belge, T., and Wolbers, T. (2023). Reaching the Goal: Superior Navigators in Late Adulthood Provide a Novel Perspective into Successful Cognitive Aging. Topics in Cognitive Science 15, 15–45. 10.1111/tops.12608.

3. Lövdén, M., Rönnlund, M., Wahlin, Å., Bäckman, L., Nyberg, L., and Nilsson, L.-G. (2004). The Extent of Stability and Change in Episodic and Semantic Memory in Old Age: Demographic Predictors of Level and Change. J Gerontol B Psychol Sci Soc Sci 59, P130–P134. 10.1093/geronb/59.3.P130.

4. Nyberg, L. (2017). Functional brain imaging of episodic memory decline in ageing. Journal of Internal Medicine 281, 65–74. 10.1111/joim.12533.

5. Hafting, T., Fyhn, M., Molden, S., Moser, M.-B., and Moser, E.I. (2005). Microstructure of a spatial map in the entorhinal cortex. Nature 436, 801–806. 10.1038/nature03721.

6. Doeller, C.F., Barry, C., and Burgess, N. (2010). Evidence for grid cells in a human memory network. Nature 463, 657–661. 10.1038/nature08704.

7. Stangl, M., Achtzehn, J., Huber, K., Dietrich, C., Tempelmann, C., and Wolbers, T. (2018). Compromised Grid-Cell-like Representations in Old Age as a Key Mechanism to Explain Age-Related Navigational Deficits. Current Biology 28, 1108–1115.e6. 10.1016/j.cub.2018.02.038.

8. Barnes, C.A., Suster, M.S., Shen, J., and McNaughton, B.L. (1997). Multistability of cognitive maps in the hippocampus of old rats. Nature 388, 272–275. 10.1038/40859.

9. Shen, J., Barnes, C.A., McNaughton, B.L., Skaggs, W.E., and Weaver, K.L. (1997). The Effect of Aging on Experience-Dependent Plasticity of Hippocampal Place Cells. J. Neurosci. 17, 6769–6782. 10.1523/JNEUROSCI.17-17-06769.1997.

10. Herber, C.S., Pratt, K.J.B., Shea, J.M., Villeda, S.A., and Giocomo, L.M. (2025). Spatial coding dysfunction and network instability in the aging medial entorhinal cortex. Nat Commun 16, 8770. 10.1038/s41467-025-63229-0.

11. Burak, Y., and Fiete, I.R. (2009). Accurate Path Integration in Continuous Attractor Network Models of Grid Cells. PLOS Computational Biology 5, e1000291. 10.1371/journal.pcbi.1000291.

12. Weber, S.N., and Sprekeler, H. (2019). A local measure of symmetry and orientation for individual spikes of grid cells. PLOS Computational Biology 15, e1006804. 10.1371/journal.pcbi.1006804.

13. Bin Khalid, I., Reifenstein, E.T., Auer, N., Kunz, L., and Kempter, R. (2024). Quantitative modeling of the emergence of macroscopic grid-like representations. eLife 13, e85742. 10.7554/eLife.85742.

14. Winter, S.S., Clark, B.J., and Taube, J.S. (2015). Spatial navigation. Disruption of the head direction cell network impairs the parahippocampal grid cell signal. Science 347, 870–874. 10.1126/science.1259591.

15. Baumann, O., and Mattingley, J.B. (2010). Medial parietal cortex encodes perceived heading direction in humans. J Neurosci 30, 12897–12901. 10.1523/JNEUROSCI.3077-10.2010.

16. Chadwick, M.J., Jolly, A.E., Amos, D.P., Hassabis, D., and Spiers, H.J. (2015). A Goal Direction Signal in the Human Entorhinal/Subicular Region. Current Biology 25, 87. 10.1016/j.cub.2014.11.001.

17. Shine, J.P., Valdés-Herrera, J.P., Tempelmann, C., and Wolbers, T. (2019). Evidence for allocentric boundary and goal direction information in the human entorhinal cortex and subiculum. Nat Commun 10, 4004. 10.1038/s41467-019-11802-9.

18. Shine, J.P., Valdés-Herrera, J.P., Hegarty, M., and Wolbers, T. (2016). The Human Retrosplenial Cortex and Thalamus Code Head Direction in a Global Reference Frame. J. Neurosci. 36, 6371–6381. 10.1523/JNEUROSCI.1268-15.2016.

19. Bernard, M., Shine, J., Bicanski, A., and Wolbers, T. (2025). How human aging disrupts the head direction network: evidence from VR experiments and mechanistic models. Preprint at bioRxiv, 10.1101/2025.11.24.690256 https://doi.org/10.1101/2025.11.24.690256.

20. Stangl, M., Shine, J., and Wolbers, T. (2017). The GridCAT: A Toolbox for Automated Analysis of Human Grid Cell Codes in fMRI. Front. Neuroinform. 11. 10.3389/fninf.2017.00047.

21. Bicanski, A., and Burgess, N. (2016). Environmental Anchoring of Head Direction in a Computational Model of Retrosplenial Cortex. J. Neurosci. 36, 11601–11618. 10.1523/JNEUROSCI.0516-16.2016.

22. Robertson, R.G., Rolls, E.T., Georges-François, P., and Panzeri, S. (1999). Head direction cells in the primate pre-subiculum. Hippocampus 9, 206–219. 10.1002/(SICI)1098-1063(1999)9:3%3C206::AID-HIPO2%3E3.0.CO;2-H.

23. Duszkiewicz, A.J., Fricker, D., Burgalossi, A., and Peyrache, A. (2025). The postsubiculum as a head-direction cortex. Trends in Neurosciences 48, 829–840. 10.1016/j.tins.2025.09.004.

24. Cho, J., and Sharp, P.E. (2001). Head direction, place, and movement correlates for cells in the rat retrosplenial cortex. Behavioral Neuroscience 115, 3–25. 10.1037/0735-7044.115.1.3.

25. Salthouse, T. (2012). Consequences of Age-Related Cognitive Declines. Annual Review of Psychology 63, 201–226. 10.1146/annurev-psych-120710-100328.

26. Li, R., Yin, S., Zhu, X., Ren, W., Yu, J., Wang, P., Zheng, Z., Niu, Y.-N., Huang, X., and Li, J. (2017). Linking Inter-Individual Variability in Functional Brain Connectivity to Cognitive Ability in Elderly Individuals. Front. Aging Neurosci. 9. 10.3389/fnagi.2017.00385.

27. Andrews-Hanna, J.R., Snyder, A.Z., Vincent, J.L., Lustig, C., Head, D., Raichle, M.E., and Buckner, R.L. (2007). Disruption of Large-Scale Brain Systems in Advanced Aging. Neuron 56, 924–935. 10.1016/j.neuron.2007.10.038.

28. Segen, V., Stangl, M., Shine, J., and Wolbers, T. (2025). Altered Coding of Environmental Boundaries in Human Aging: An fMRI Study. Hippocampus 35, e70044. 10.1002/hipo.70044.

29. McQuail, J.A., Barnes, C.A., Bizon, J.L., Burke, S.N., Disterhoft, J.F., Foster, T.C., Huentelman, M., Kaczorowski, C.C., Long, J.M., Moore, S.J., et al. (2027). Individual Differences in Cognitive Aging Rodent Datasets (ID-CARD): A collaborative platform for behavioral analysis across the lifespan. Neurobiology of Aging 169, 27–36. 10.1016/j.neurobiolaging.2026.09.003.

30. Yoon, K., Buice, M.A., Barry, C., Hayman, R., Burgess, N., and Fiete, I.R. (2013). Specific evidence of low-dimensional continuous attractor dynamics in grid cells. Nat Neurosci 16, 1077–1084. 10.1038/nn.3450.

31. Sit, K.K., and Goard, M.J. (2023). Coregistration of heading to visual cues in retrosplenial cortex. Nat Commun 14, 1992. 10.1038/s41467-023-37704-5.

32. Marchette, S.A., Vass, L.K., Ryan, J., and Epstein, R.A. (2014). Anchoring the neural compass: coding of local spatial reference frames in human medial parietal lobe. Nat Neurosci 17, 1598–1606. 10.1038/nn.3834.

33. Taube, J.S. (2007). The Head Direction Signal: Origins and Sensory-Motor Integration. Annual Review of Neuroscience 30, 181–207. 10.1146/annurev.neuro.29.051605.112854.

34. Kononenko, N.L., and Witter, M.P. (2012). Presubiculum layer III conveys retrosplenial input to the medial entorhinal cortex. Hippocampus 22, 881–895. 10.1002/hipo.20949.

35. Insausti, R., Amaral, D.G., and Cowan, W.M. (1987). The entorhinal cortex of the monkey: II. Cortical afferents. Journal of Comparative Neurology 264, 356–395. 10.1002/cne.902640306.

36. Dubanet, O., and Higley, M.J. (2024). Retrosplenial inputs drive visual representations in the medial entorhinal cortex. Cell Reports 43. 10.1016/j.celrep.2024.114470.

37. Hu, X.-Q., Wu, K.L.-K., Rong, K.-L., Ke, Y., Yung, W.-H., Shum, D.K.-Y., and Chan, Y.-S. (2025). Harnessing a noncanonical vestibular input in the head-direction network to rectify age-related navigational deficits. Nat Aging 5, 1079–1096. 10.1038/s43587-025-00884-4.

38. Jiang, S., Hijazi, S., Sárkány, B., Gautsch, V.G., LaChance, P.A., Hasselmo, M.E., Bannerman, D., and Viney, T.J. (2025). Pathological tau alters head direction signaling and induces spatial disorientation. Cell Reports 44. 10.1016/j.celrep.2025.116610.

39. Chaudhuri, R., Gerçek, B., Pandey, B., Peyrache, A., and Fiete, I. (2019). The intrinsic attractor manifold and population dynamics of a canonical cognitive circuit across waking and sleep. Nat Neurosci 22, 1512–1520. 10.1038/s41593-019-0460-x.

40. Ajabi, Z., Keinath, A.T., Wei, X.-X., and Brandon, M.P. (2023). Population dynamics of head-direction neurons during drift and reorientation. Nature 615, 892–899. 10.1038/s41586-023-05813-2.

41. Hardcastle, K., Ganguli, S., and Giocomo, L.M. (2015). Environmental Boundaries as an Error Correction Mechanism for Grid Cells. Neuron 86, 827–839. 10.1016/j.neuron.2015.03.039.

42. Segen, V., Ying, J., Morgan, E., Brandon, M., and Wolbers, T. (2022). Path integration in normal aging and Alzheimer’s disease. Trends in Cognitive Sciences 26, 142–158. 10.1016/j.tics.2021.11.001.

43. Naveilhan, C., Sicard, M., Zory, R., Gramann, K., and Ramanoël, S. (2026). The role of cognitive-motor interaction in landmark reliance and navigational deficits in older adults. Neurobiology of Aging 168, 109–122. 10.1016/j.neurobiolaging.2026.07.011.

44. Sugar, J., and Moser, M.-B. (2019). Episodic memory: Neuronal codes for what, where, and when. Hippocampus 29, 1190–1205. 10.1002/hipo.23132.

45. Dong, L.L., and Fiete, I.R. (2024). Grid Cells in Cognition: Mechanisms and Function. Annual Review of Neuroscience 47, 345–368. 10.1146/annurev-neuro-101323-112047.

46. Leventhal, A.G., Wang, Y., Pu, M., Zhou, Y., and Ma, Y. (2003). GABA and Its Agonists Improved Visual Cortical Function in Senescent Monkeys. Science 300, 812–815. 10.1126/science.1082874.

47. Bürkner, P.-C. (2017). brms: An R Package for Bayesian Multilevel Models Using Stan. Journal of Statistical Software 80, 1–28. 10.18637/jss.v080.i01.

48. Yushkevich, P.A., Amaral, R.S.C., Augustinack, J.C., Bender, A.R., Bernstein, J.D., Boccardi, M., Bocchetta, M., Burggren, A.C., Carr, V.A., Chakravarty, M.M., et al. (2015). Quantitative comparison of 21 protocols for labeling hippocampal subfields and parahippocampal subregions in in vivo MRI: towards a harmonized segmentation protocxol. Neuroimage 111, 526–541. 10.1016/j.neuroimage.2015.01.004.

49. Mazloum-Farzaghi, N., Barense, M.D., Ryan, J.D., Stark, C.E.L., and Olsen, R.K. (2024). The Effect of Segmentation Method on Medial Temporal Lobe Subregion Volumes in Aging. Human Brain Mapping 45, e70054. 10.1002/hbm.70054.

50. Tournier, J.-D., Smith, R., Raffelt, D., Tabbara, R., Dhollander, T., Pietsch, M., Christiaens, D., Jeurissen, B., Yeh, C.-H., and Connelly, A. (2019). MRtrix3: A fast, flexible and open software framework for medical image processing and visualisation. Neuroimage 202, 116137. 10.1016/j.neuroimage.2019.116137.

51. Garyfallidis, E., Brett, M., Amirbekian, B., Rokem, A., Van Der Walt, S., Descoteaux, M., and Nimmo-Smith, I. (2014). Dipy, a library for the analysis of diffusion MRI data. Front. Neuroinform. 8. 10.3389/fninf.2014.00008.

52. Tustison, N.J., Cook, P.A., Holbrook, A.J., Johnson, H.J., Muschelli, J., Devenyi, G.A., Duda, J.T., Das, S.R., Cullen, N.C., Gillen, D.L., et al. (2021). The ANTsX ecosystem for quantitative biological and medical imaging. Sci Rep 11, 9068. 10.1038/s41598-021-87564-6.

53. Behrenbruch, N., Schwarck, S., Schumann-Werner, B., Molloy, E.N., Garcia-Garcia, B., Hochkeppler, A., Fischer, L., Büchel, A.-T., Incesoy, E.I., Bernal, J., et al. (2026). A physically and mentally active lifestyle relates to younger brain and cognitive age. GeroScience 48, 1853–1873. 10.1007/s11357-025-01764-w.

54. Schmid, N.S., Ehrensperger, M.M., Berres, M., Beck, I.R., and Monsch, A.U. (2014). The Extension of the German CERAD Neuropsychological Assessment Battery with Tests Assessing Subcortical, Executive and Frontal Functions Improves Accuracy in Dementia Diagnosis. Dement Geriatr Cogn Disord Extra 4, 322–334. 10.1159/000357774.

55. Nasreddine, Z.S., Phillips, N.A., Bédirian, V., Charbonneau, S., Whitehead, V., Collin, I., Cummings, J.L., and Chertkow, H. (2005). The Montreal Cognitive Assessment, MoCA: A brief screening tool for mild cognitive impairment. Journal of the American Geriatrics Society 53, 695–699. 10.1111/j.1532-5415.2005.53221.x.

56. Carson, N., Leach, L., and Murphy, K.J. (2018). A re-examination of Montreal Cognitive Assessment (MoCA) cutoff scores. International Journal of Geriatric Psychiatry 33, 379–388. 10.1002/gps.4756.

57. ASHS Documentation https://sites.google.com/view/ashs-dox/.

58. Olsen, R.K., Yeung, L.-K., Noly-Gandon, A., D’Angelo, M.C., Kacollja, A., Smith, V.M., Ryan, J.D., and Barense, M.D. (2017). Human anterolateral entorhinal cortex volumes are associated with cognitive decline in aging prior to clinical diagnosis. Neurobiology of Aging 57, 195–205. 10.1016/j.neurobiolaging.2017.04.025.

59. Navarro Schröder, T., Haak, K.V., Zaragoza Jimenez, N.I., Beckmann, C.F., Doeller, C.F. (2015). Functional topography of the human entorhinal cortex. eLife 4, e06738. 10.7554/eLife.06738.

60. Kunz, L., Schröder, T.N., Lee, H., Montag, C., Lachmann, B., Sariyska, R., Reuter, M., Stirnberg, R., Stöcker, T., Messing-Floeter, P.C., et al. (2015). Reduced grid-cell–like representations in adults at genetic risk for Alzheimer’s disease. Science 350, 430–433. 10.1126/science.aac8128.

61. Bierbrauer, A., Kunz, L., Gomes, C.A., Luhmann, M., Deuker, L., Getzmann, S., Wascher, E., Gajewski, P.D., Hengstler, J.G., Fernandez-Alvarez, M., et al. (2020). Unmasking selective path integration deficits in Alzheimer’s disease risk carriers. Science Advances 6, eaba1394. 10.1126/sciadv.aba1394.

62. Liang, Z., Wu, S., Wu, J., Wang, W.-X., Qin, S., and Liu, C. (2024). Distance and grid-like codes support the navigation of abstract social space in the human brain. eLife 12, RP89025. 10.7554/eLife.89025.

63. Dalton, M.A., Zeidman, P., Barry, D.N., Williams, E., and Maguire, E.A. (2017). Segmenting subregions of the human hippocampus on structural magnetic resonance image scans: An illustrated tutorial. Brain and Neuroscience Advances 1, 2398212817701448. 10.1177/2398212817701448.

64. Desikan, R.S., Ségonne, F., Fischl, B., Quinn, B.T., Dickerson, B.C., Blacker, D., Buckner, R.L., Dale, A.M., Maguire, R.P., Hyman, B.T., et al. (2006). An automated labeling system for subdividing the human cerebral cortex on MRI scans into gyral based regions of interest. NeuroImage 31, 968–980. 10.1016/j.neuroimage.2006.01.021.

65. SPM12 Software - Statistical Parametric Mapping https://www.fil.ion.ucl.ac.uk/spm/software/spm12/.

66. Behzadi, Y., Restom, K., Liau, J., and Liu, T.T. (2007). A component based noise correction method (CompCor) for BOLD and perfusion based fMRI. NeuroImage 37, 90–101. 10.1016/j.neuroimage.2007.04.042.

67. Grydeland, H., Walhovd, K.B., Tamnes, C.K., Westlye, L.T., and Fjell, A.M. (2013). Intracortical Myelin Links with Performance Variability across the Human Lifespan: Results from T1- and T2-Weighted MRI Myelin Mapping and Diffusion Tensor Imaging. J. Neurosci. 33, 18618–18630. 10.1523/JNEUROSCI.2811-13.2013.

