## Supplemental materials for "Temporal instability weakens entorhinal grid-like coding in older adults"

**Table S1. Bayesian regression results for the off-diagonal hexadirectional-signal sensitivity analysis**

| Hemisphere | Predictor | β | Posterior SD | 95% CrI | *pd* |
|---|---|---|---|---|---|
| **Left pmEC** | **Hexadirectional orientation stability** | **0.267** | **0.107** | **[0.054, 0.475]** | **.993** |
| **Left pmEC** | Hexadirectional geometry stability | 0.029 | 0.108 | [−0.186, 0.240] | .609 |
| **Left pmEC** | Hexadirectional spatial stability | 0.059 | 0.109 | [−0.156, 0.273] | .706 |
| **Left pmEC** | Age | 0.084 | 0.110 | [−0.130, 0.297] | .779 |
| **Right pmEC** | Hexadirectional orientation stability | −0.133 | 0.108 | [−0.345, 0.077] | .108 |
| **Right pmEC** | Hexadirectional geometry stability | 0.211 | 0.108 | [−0.003, 0.423] | .973 |
| **Right pmEC** | Hexadirectional spatial stability | −0.026 | 0.107 | [−0.240, 0.186] | .400 |
| **Right pmEC** | Age | −0.022 | 0.108 | [−0.234, 0.190] | .415 |

*Note*; All models used (n = 85), pd is the posterior probability that the coefficient is positive. CrI, credible interval; pmEC, postetiot-medial-entorhinal-cortex

**Supplementary Table S2. Associations of pmEC hexadirectional signal with object-location accuracy**

| Hemisphere | Predictor | Log-odds β | 95% CrI |
|---|---|---|---|
| **Left pmEC** | Hexadirectional signal | 0.122 | [−0.017, 0.261] |
| **Left pmEC** | Age | −0.315 | [−0.457, −0.174] |
| **Right pmEC** | Hexadirectional signal | 0.136 | [−0.009, 0.277] |
| **Right pmEC** | Age | −0.300 | [−0.439, −0.160] |

*Note.* Results are from separate age-adjusted Bayesian binomial models predicting correct responses on the object-location task relative to trials answered ($n$ = 83). Continuous predictors were z-standardized; coefficients are on the log-odds scale.

**Supplementary Table S3. Associations between right-pmEC structural properties and right-pmEC hexadirectional signal and hexadirectional orientation stability**

| Outcome | Structural predictor | *n* | Posterior β | Posterior SD | 95% CrI | *pd* | Median $BF_{01}$ |
|---|---|---|---|---|---|---|---|
| **Right-pmEC hexadirectional signal** | Mean diffusivity | 61 | 0.129 | 0.131 | [−0.129, 0.386] | .840 | 2.31 |
| **Right-pmEC hexadirectional signal** | T1w/T2w ratio | 85 | −0.047 | 0.108 | [−0.260, 0.167] | .671 | **4.18** |
| **Right-pmEC hexadirectional signal** | TIV-residualised volume | 85 | −0.152 | 0.107 | [−0.364, 0.058] | .923 | 1.67 |
| **Right-pmEC hexadirectional orientation stability** | Mean diffusivity | 61 | −0.085 | 0.130 | [−0.342, 0.172] | .744 | **3.09** |
| **Right-pmEC hexadirectional orientation stability** | T1w/T2w ratio | 85 | −0.091 | 0.109 | [−0.305, 0.125] | .798 | **3.24** |
| **Right-pmEC hexadirectional orientation stability** | TIV-residualised volume | 85 | 0.057 | 0.109 | [−0.156, 0.271] | .701 | **4.05** |

*Note.* All models included age as a covariate and used standardised variables. $BF_{01}$ compares the age-only model with the model additionally including the structural predictor; values above 1 favour the age-only model. *pd*, probability of direction, denotes the posterior probability of the direction shown by the estimate. CrI, credible interval; pmEC, postetiot-medial-entorhinal-cortex; TIV, total intracranial volume.

**Supplementary Table S4. Bayesian mediation analysis of retrosplenial absolute-orientation stability, left-pmEC hexadirectional orientation stability, and left-pmEC hexadirectional signal**

| Effect | Path | Posterior estimate | Posterior SD | 95% CrI | *pd* |
|---|---|---|---|---|---|
| ***a* path** | Retrosplenial absolute-orientation stability → left-pmEC hexadirectional orientation stability | 0.222 | 0.107 | [0.014, 0.433] | .982 |
| ***b* path** | Left-pmEC hexadirectional orientation stability → left-pmEC hexadirectional signal | 0.562 | 0.094 | [0.377, 0.746] | > .999 |
| **Indirect effect** | Retrosplenial absolute-orientation stability → left-pmEC hexadirectional orientation stability → left-pmEC hexadirectional signal | 0.125 | 0.064 | [0.008, 0.261] | .982 |
| **Direct effect (*c′*)** | Retrosplenial absolute-orientation stability → left-pmEC hexadirectional signal, conditional on left-pmEC hexadirectional orientation stability | −0.059 | 0.093 | [−0.245, 0.123] | .740 |
| **Total effect (*c*)** | Retrosplenial absolute-orientation stability → left-pmEC hexadirectional signal | 0.066 | 0.109 | [−0.147, 0.281] | .727 |

*Note.* RSC, retrosplenial cortex; pmEC, posterior-medial-entorhinal-cortex; CrI, credible interval; *pd*, probability of direction. All variables were standardised and models included age as a covariate. $N = 85$. The *b* and direct-effect paths were estimated jointly in the outcome model. For negative estimates, *pd* denotes the posterior probability of a negative effect.
